# Ionic polyphosphorylation of histidine repeat proteins by inorganic polyphosphate

**DOI:** 10.1101/2023.04.10.536149

**Authors:** Nolan Neville, Kirsten Lehotsky, Zhiyun Yang, Kody A. Klupt, Alix Denoncourt, Michael Downey, Zongchao Jia

**Affiliations:** Department of Biomedical and Molecular Sciences, Queen’s University, Kingston, ON K7L 3N6, Canada; Department of Cellular and Molecular Medicine, University of Ottawa, Ottawa, ON K1H 8M5, Canada; Ottawa Institute of Systems Biology, Ottawa, ON K1H 8M5, Canada

## Abstract

Inorganic polyphosphate (polyP) is a linear polymer of orthophosphate that is present in nearly all organisms studied to date. A remarkable function of polyP involves its attachment to lysine residues via non-enzymatic post-translational modification (PTM) that is presumed to be covalent. Here, we show that proteins containing tracts of consecutive histidine residues exhibit a similar modification by polyP, which confers an electrophoretic mobility shift on NuPAGE gels. Our screen uncovered 30 human and yeast histidine repeat proteins that are specifically modified by polyP. This polyP modification is histidine-dependent and non-covalent in nature, though remarkably, it withstands harsh denaturing conditions—a hallmark of covalent PTMs. We have termed this interaction ionic histidine polyphosphorylation (iH-PPn) to describe its unique PTM-like properties. Importantly, we show that iH-PPn disrupts phase separation and phosphorylation activity of the human protein kinase DYRK1A, and inhibits the activity of the transcription factor MafB, highlighting iH-PPn as a potential hitherto unrecognized regulatory mechanism.

**GRAPHICAL ABSTRACT.**
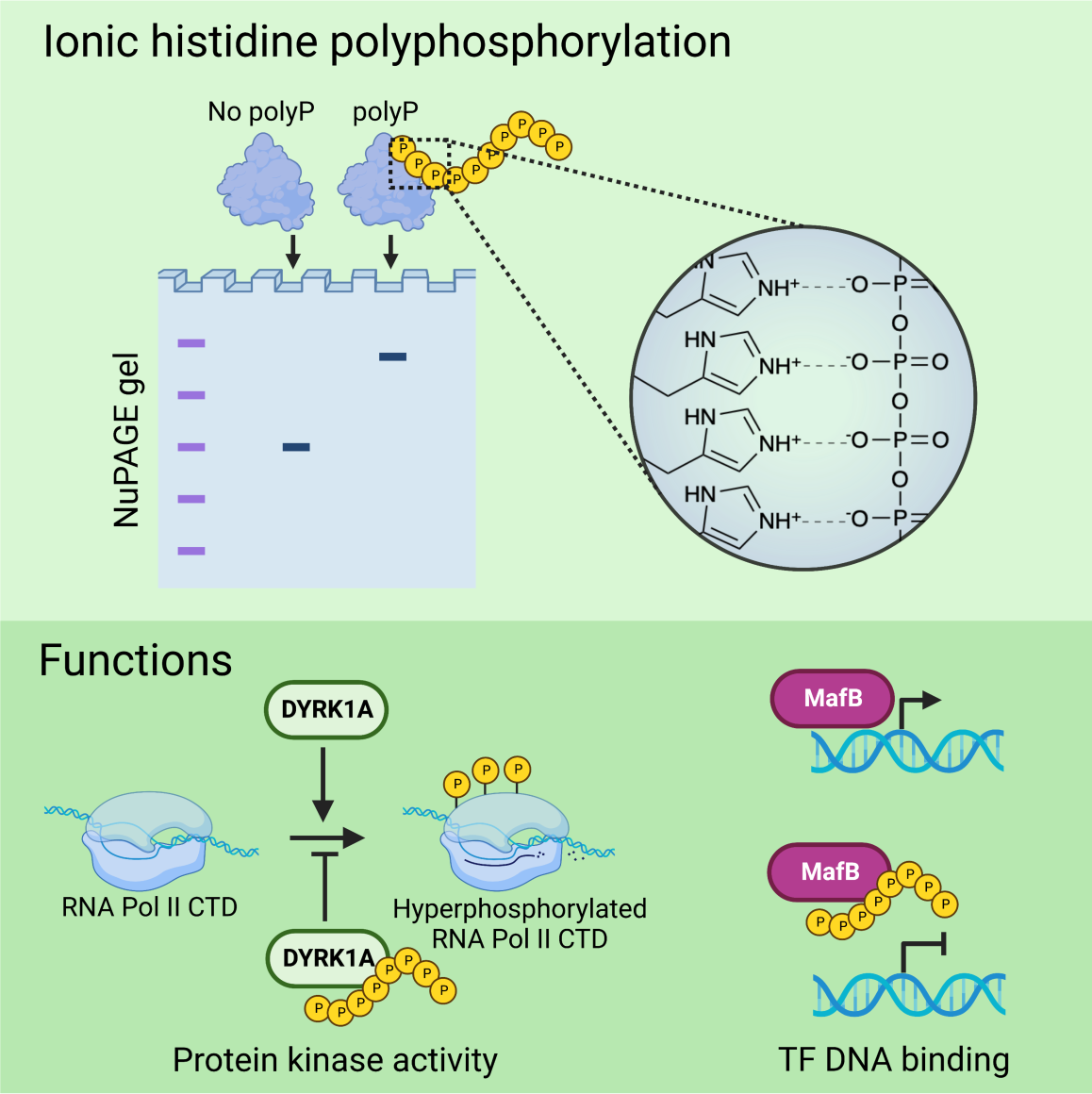

## HIGHLIGHTS

- Identified 20 human and 10 yeast histidine repeat proteins that are modified by polyphosphorylation
- Histidine repeat sequences are the direct target of polyP modification
- The polyP-histidine modification is non-covalent but resistant to extreme denaturing conditions
- Histidine polyphosphorylation negatively regulates the activities of protein kinase DYRK1A and transcription factor MafB

## INTRODUCTION

Inorganic polyphosphate (polyP) is a chain of phosphate residues ranging from three to thousands of units in length. PolyP has been detected in all domains of life, ranging from bacteria to human cells.^1^ In bacteria, polyP is synthesized by polyP kinases PPK1 and PPK2,^2,3^ and polyP levels can reach upwards of 50 mM in *Escherichia coli*.^4,5^ In the budding yeast *Saccharomyces cerevisiae*, polyP is synthesized by the vacuolar transporter chaperone (VTC) complex^6^ and can accumulate to >200 mM.^7^ The source of polyP in higher eukaryotes is still unclear, although a role for the F_0_F_1_ ATPase has been proposed.^8^ Mammals encode no obvious homologues of bacterial PPK or yeast VTC enzymes. Mammalian polyP levels are more typically on the micromolar scale, ranging from 25 to 120 µM in rodent cell homogenates,^9^ though human platelets accumulate up to 130 mM polyP in dense granules.^10^ Remarkably, polyP can be attached to proteins via what is postulated to be a non-enzymatic post-translational modification (PTM) termed lysine polyphosphorylation (K-PPn).^11^ Azevedo *et al.* discovered that the yeast proteins Top1 and Nsr1 undergo K-PPn, resulting in a dramatic decrease in mobility during electrophoresis on NuPAGE that led the authors to posit a covalent attachment.^12^ K-PPn disrupted the localization and protein-protein interaction of Nsr1 and Top1 and inhibited the *in vitro* topoisomerase activity of Top1. The lysine residues targeted by polyP in each protein were mapped to polyacidic serine and lysine rich (PASK) clusters.^12^ Subsequent bioinformatic screening for other proteins that contain the PASK motif—defined as a stretch of 20 amino acids with at least 75 % D/E/S and at least one K— identified an additional 23 hits from yeast and 6 human proteins that undergo polyphosphorylation.^13,14^ A microarray screen added 8 more human proteins that exhibit a polyP-dependent NuPAGE shift, though 4 of these hits contain no obvious PASK motif.^15,16^

PolyP also interacts non-covalently with numerous proteins. For example, proteins containing a conserved histidine α-helical domain (CHAD) bind to polyP via two 4-helical bundles with a highly basic central cavity.^17^ PolyP in human blood is strongly procoagulant, and several proteins involved in blood clotting have been shown to interact with polyP.^18–20^ Histidine-rich glycoprotein (HRG) is a plasma protein with anticoagulant properties and an unusually high composition (12.6 %) of histidine residues. HRG binds polyP with *K*_d_ ∼ 152 nM in a Zn^2+^-dependent manner to attenuate its procoagulant effects.^21^ A screen of 15 000 human proteins yielded 309 polyP binding hits, though only five of these were validated to interact with polyP electrostatically.^15^ PolyP also functions as a primordial chaperone, interacting with a diverse range of client proteins to maintain them in a soluble state.^5^

Single amino acid repeats (SARs) are common in eukaryotes and are present in 18-20% of human proteins.^22^ The human genome encodes 86 proteins with five or more consecutive histidine residues.^23^ Histidine repeats have been ascribed a variety of functions, including driving liquid-liquid phase separation^24^ and protein-protein interactions.^25^ Histidine repeats are overrepresented in nuclear proteins, especially those involved in neurodevelopment, and histidine repeats are both necessary and sufficient to target proteins to nuclear speckles.^23^

We discovered that proteins containing stretches of consecutive histidine residues are modified by polyP, as observed via electrophoretic mobility shift on NuPAGE gels. We show that histidine residues mediate ionic polyP binding to these proteins, which we have therefore termed ionic histidine polyphosphorylation (iH-PPn). This modification alters the activity of the clinically relevant human histidine repeat proteins DYRK1A and MafB, suggesting iH-PPn as a global protein modification with potentially widespread regulatory implications.

## RESULTS

### Histidine repeats confer a polyP dependent NuPAGE shift

During our previous studies of His-tagged PPK1 from *Pseudomonas aeruginosa,*^26^ we observed an electrophoretic mobility shift in the presence of polyP on NuPAGE gels as previously documented for K-PPn.^13^ Removal of the hexahistidine tag eliminated the NuPAGE shift, suggesting the histidine repeat as the target of polyP modification (Figures S1A-S1C).

To test if proteins containing endogenous stretches of consecutive histidine also exhibited a polyP-dependent NuPAGE shift, we selected 27 human proteins with five or more consecutive histidine residues (Figure 1A). Selected proteins were expressed in *E. coli* with N-terminal maltose binding protein (MBP) tags. Of the 27 candidates, 19 proteins displayed a polyP-dependent NuPAGE shift (Figures 1B and 1C; Figure S1E). MBP alone and eight of the candidate fusions displayed no change in mobility.

**Figure 1.**
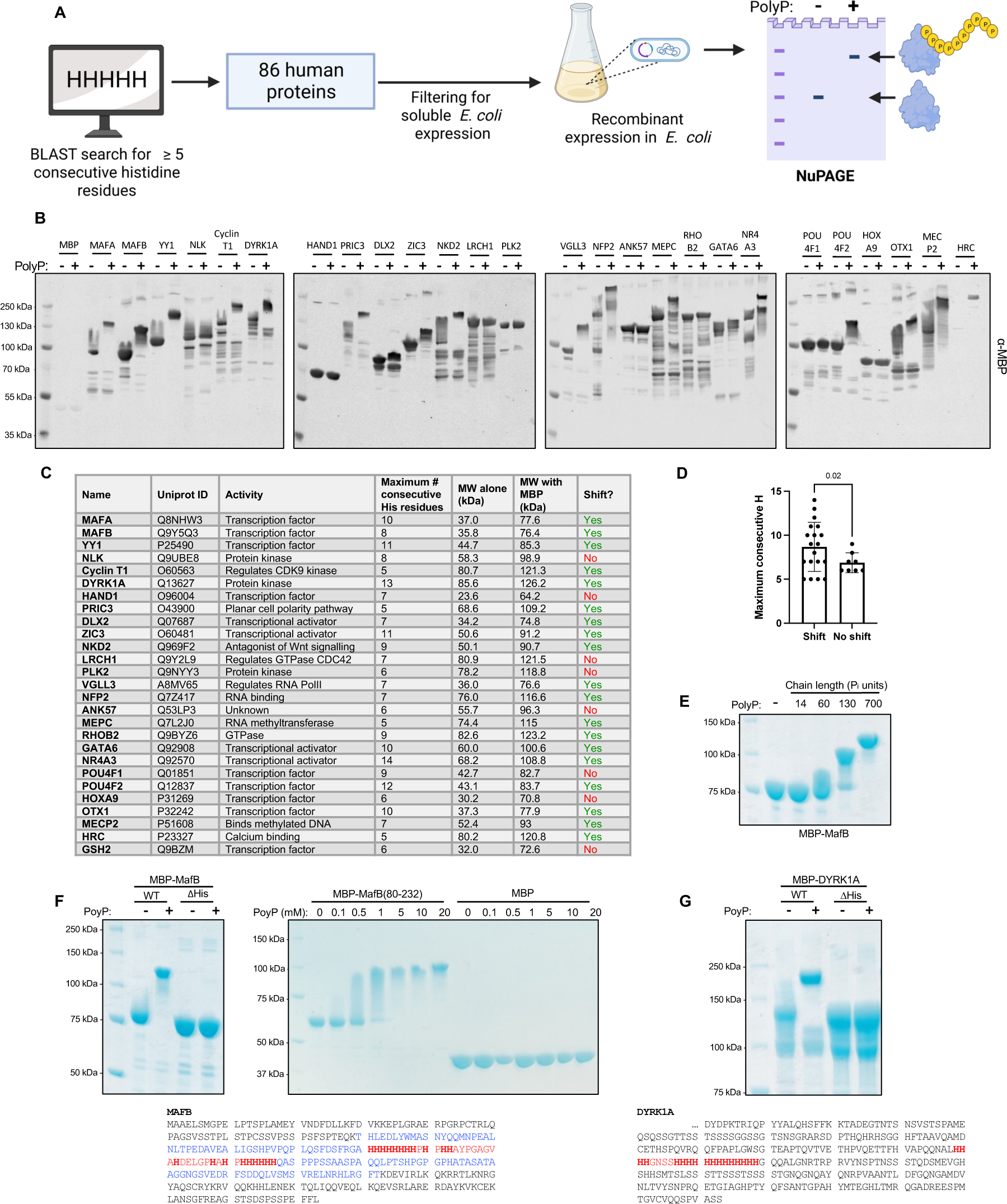
Identification of polyphosphorylated human histidine repeat proteins. (A) Parameters used to select human histidine repeat candidates (B) NuPAGE analysis of histidine repeat candidates. (C) Summary of proteins screened. (D) NuPAGE shift versus histidine repeat length (unpaired t-test with Welch’s correction). (E) NuPAGE shift is polyP chain length-dependent. (F and G) Deleting the histidine repeat region of MafB (red; residues 131-167) or DYRK1A (red; residues 599-619) eliminates NuPAGE shift. MafB histidine repeat region (blue; residues 80-232) confers a dose-dependent polyP shift to MBP.

Likelihood of shift correlated with the length of the histidine repeat, with an average of 8.7 consecutive H in shifted proteins versus 6.9 consecutive H in non-shifted proteins (Figure 1D). Screening of 24 histidine repeat proteins from *S. cerevisiae* yielded 10 additional hits including the important glucose-regulated kinase Snf1 (Figure S1D; Table S1).

Many of the hit proteins are known to play critical roles in human physiology (Figure 1C). For example, MafA and MafB are transcription factors that regulate insulin gene expression and hematopoiesis, respectively.^27,28^ Dual-specificity tyrosine phosphorylation-regulated kinase 1A (DYRK1A) directly phosphorylates RNA polymerase II to activate transcription,^29^ and DYRK1A has been implicated in neurodevelopment and is overexpressed in Down syndrome.^30^ Given their relatively well-characterized biological functions, we selected MafB, DYRK1A, and Snf1 as model proteins to further define the polyP binding interaction. Treatment of MBP-MafB with polyP of varying chain lengths revealed that its NuPAGE shift is polyP length-dependent (Figure 1E). Deletion of the histidine repeat region in MafB (residues 131-167) and DYRK1A (residues 599-619) abolished the shift for both proteins (Figures 1F and 1G).

To exclude the possibility of lysine polyphosphorylation, we fused the largest possible stretch of residues flanking the histidine-rich region of MafB that contained no lysine residues (80-232) to MBP, which conferred a polyP concentration-dependent shift (Figure 1F). To exclude the possibility of bacterial expression artifacts influencing this modification, we immunoblotted for endogenous cyclin T1 and YY1 in HeLa lysate treated with polyP and observed shifts similar to those of the recombinant proteins (Figures S1F and S1G).

### The histidine-polyP interaction is non-covalent, but strongly ionic and specific

The fact that the polyP-dependent histidine shift was observed even after samples were boiled in denaturing buffer and run on denaturing NuPAGE gels led us to assume this modification was covalent, as is presumed to be the case for K-PPn.^12^ Moreover, addition of tetramethylethylenediamine (TEMED) to homemade Bis-Tris gels collapsed the shift (Figure S2A), as previously observed for K-PPn.^13^ To verify that the NuPAGE shifting was not merely an artifact of electrophoresis, we used size exclusion chromatography (SEC) to monitor the interaction between proteins and polyP in solution. MBP alone exhibited no shift in elution volume upon polyP treatment, whereas MBP fused to the histidine repeat domain (residues 1-65) of Snf1 (Figure 2F) eluted significantly earlier in the presence of polyP (Figure 2A), mirroring the NuPAGE shift results. Surprisingly, increasing the pH above 7.5 disrupted protein binding to polyP as observed via SEC and polyP pulldown (Figures 2B and 2C). This trend was also observed for MBP-MafB(80-232) (Figures S2B-S2C). Increasing the ionic strength (NaCl or (NH_4_)_2_SO_4_) of the buffer also disrupted the interaction, and this was much more pronounced at pH 7.5 versus pH 6.2 (Figures 2D, 2E, S2D and S2E). At pH 7.5, 50 mM NaCl abolished the shift, while even 500 mM NaCl failed to completely abolish the shift at pH 6.2. By contrast, the PASK-containing positive control MBP-Rts1(1-65) maintained a SEC elution shift at pH 8 with 100 mM NaCl (Figure S2F). Together, these results strongly suggest that the histidine-polyP interaction is ionic, rather than covalent, hence our designation as ionic histidine polyphosphorylation (iH-PPn).

**Figure 2.**
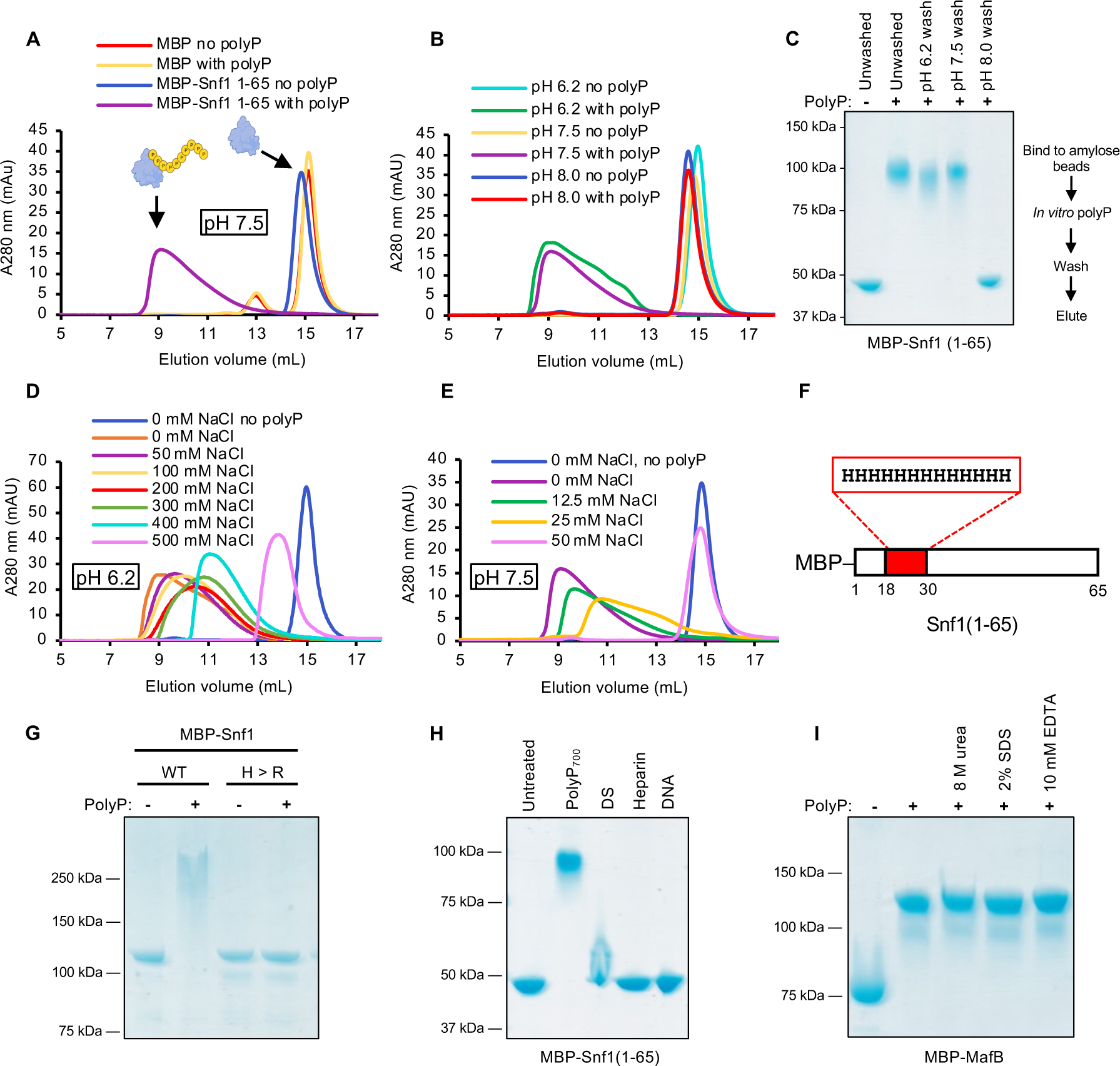
Histidine repeat polyphosphorylation is an ionic non-covalent interaction. (A) SEC of MBP-Snf1(1-65) fusion or MBP alone incubated with 5 mM polyP_700_ or 5 mM monomeric NaH_2_PO_4_. (B) SEC of MBP-Snf1(1-65) in buffers of indicated pH. (C) PolyP pulldown by MBP-Snf1(1-65) at different pH values. (D and E) MBP-Snf1(1-65) resolved by SEC in buffer supplemented with the indicated concentration of salt at pH 6.2 and 7.5, respectively. (F) Schematic of the MBP-Snf1(1-65) construct. (G) Arginine replacement NuPAGE test. (H) Polyanion NuPAGE test. DS; dextran sulfate. (I) Denaturant and chelator NuPAGE test.

The apparent ionic nature of histidine-polyP binding led us to question whether this interaction was specific, or merely an indiscriminate electrostatic attraction. To test this, we replaced the 13H tract in Snf1 with 13 arginine residues. The arginine repeat Snf1 did not shift in the presence of polyP (Figure 2G), indicating that positive charge alone is insufficient for modification by polyP. Similarly, MBP-Snf1(1-65) was largely unaffected by biological polyanions other than polyP. Heparin and single-stranded DNA had no effect on NuPAGE mobility, and dextran sulfate yielded only a faint smear upwards (Figures 2H and S3A). The histidine-polyP shift was unaffected by strong denaturants such as 8M urea or the chelator EDTA (Figure 2I).

Systematic alanine substitution within the Snf1 His repeat abolished shifting once three or fewer consecutive H remained (Figures S3B-S3C). Interestingly, HRG—which contains 12.6% histidine but never more than two consecutive H—exhibited a NuPAGE iH-PPn shift that was unaffected by EDTA or zinc (Figures S3D-S3F). *In silico* docking of a 13H peptide to polyP_13_ yielded a model in which the histidine sidechains preferentially orient to one side of the peptide to wrap around polyP in a zipper-like fashion. The predicted p*K*as of sidechains in contact with polyP are markedly higher those in the absence of polyP (Figures S3G-S3H).

### Histidine repeat alone is sufficient for iH-PPn

We explored the minimal sequence required for iH-PPn with chemically synthesized peptides tagged with fluorescein isothiocyanate (FITC). Poly-dependent shifts were observed for the 13H and the PASK peptide consisting of 17 residues from the predicted PASK motif of Rts1.^13^ Thus, consecutive histidine residues alone are sufficient for iH-PPn. By contrast, no shift was observed for the negative control peptide consisting of 13 consecutive arginine residues (Figures 3A and 3B). PolyP also blocked the ability of the 13H peptide to bind to nickel resin (Figure S3I). These trends were supported by circular dichroism (CD). 13H, PASK, and 13R peptides displayed disordered random coil conformations in the absence of polyP. In the presence of polyP, the spectra of 13H and PASK peptides shifted to more ordered conformations, as evidenced by the inflection around 180-200 nm. By contrast, the overall shape of the 13R spectrum is unaltered by polyP (Figure 3C).

**Figure 3.**
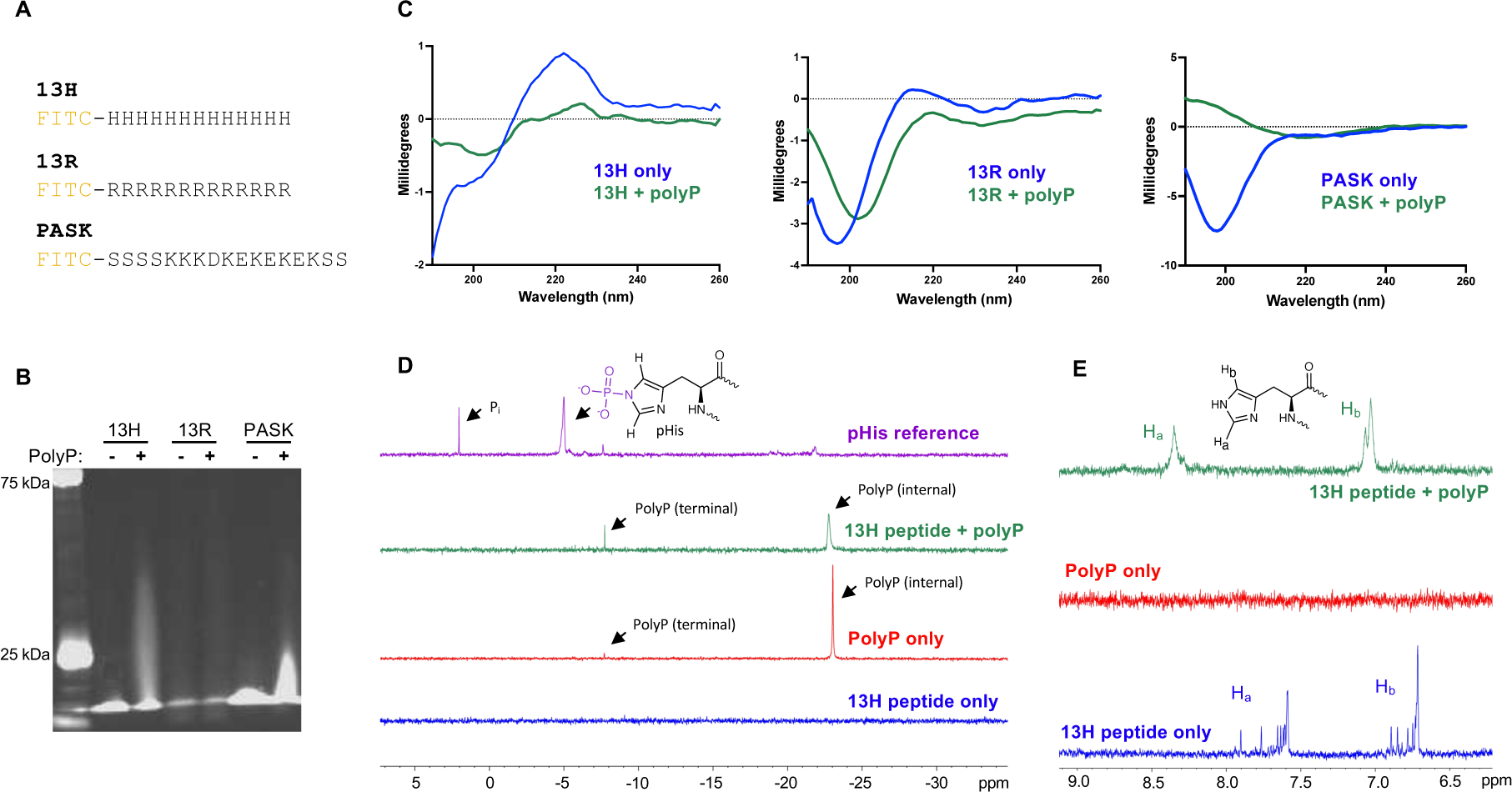
Histidine repeat peptide is sufficient for iH-PPn. (A) FITC-tagged 13H, 13R, and PASK peptide sequences. (B) NuPAGE of peptides with 5 mM polyP_700_ imaged under UV light. (C) Circular dichroism spectra of 13H, 13R and PASK peptides in the presence of 2.5 mM polyP_700_. (D) ^31^P NMR of 13H peptide with and without 5 mM polyP_700_, and a positive control phosphohistidine reference (structure inset). (E) ^1^H NMR of 13H peptide treated with 5 mM polyP_700_. Histidine sidechain proton regions labeled according to inset structure.

Having demonstrated that 13H peptide is sufficient to undergo iH-PPn, we used this peptide for NMR studies. Phosphohistidine (pHis) is known to exhibit a characteristic P-N phosphoramidate bond peak in ^31^P NMR at approximately −5 ppm,^31,32^ which we would expect to observe if polyP attachment to histidine was covalent as it is presumed to be for K-PPn. A clear phosphoramidate peak at −5 ppm is visible for the pHis positive control,^33^ but the 13H peptide treated with polyP displays no such peak (Figure 3D). The peak at ∼ −7.5 ppm, corresponding to terminal polyP phosphate groups,^34^ becomes more pronounced in the presence of peptide. Conversely, the peak at ∼ −23 ppm which corresponds to internal polyP residues^34^ becomes broadened and weaker in the presence of peptide. In ^1^H NMR, the peaks corresponding to the histidine sidechain protons are sharp but become muddled and shifted downfield in the presence of polyP (Figure 3E). The changes in ^31^P polyP peaks and ^1^H histidine peaks provide direct evidence of a polyP-histidine interaction, but the absence of a new phosphoramidate ^31^P peak is further evidence that this interaction is non-covalent.

### iH-PPn alters DYRK1A phase separation and protein kinase activity

Given that polyP binds to—and thereby likely occludes—the DYRK1A histidine repeat, we reasoned that iH-PPn might replicate the phenotypes associated with histidine repeat deletion. Histidine deletion has been shown to disrupt the ability of DYRK1A to localize to nuclear speckles, which are phase-separated compartments enriched with transcription and splicing factors.^23,35^ We therefore monitored the effect of polyphosphorylation on EGFP-tagged DYRK1A nuclear speckle formation in HeLa cells (Figure 4A). Since polyP levels in human cells are usually low, HeLa cells were co-transfected with a plasmid expressing mCherry fused to *P. aeruginosa* PPK1 to drive overproduction of polyP, the success of which was confirmed by the NuPAGE shift of WT EGFP-DYRK1 in the lysate of co-transfected cells and by direct detection of polyP via negative DAPI staining (Figures S4A-S4B). WT EGFP-DYRK1A formed bright puncta indicative of nuclear speckles when co-transfected with plasmid expressing mCherry alone, while co-transfection with the PPK1-mCherry fusion abolished the EGFP puncta resulting in diffuse green fluorescence in the nucleoplasm. EGFP-ΔHis DYRK1A was unable to form nuclear speckles in the presence or absence of PPK1 (Figure 4A).

**Figure 4.**
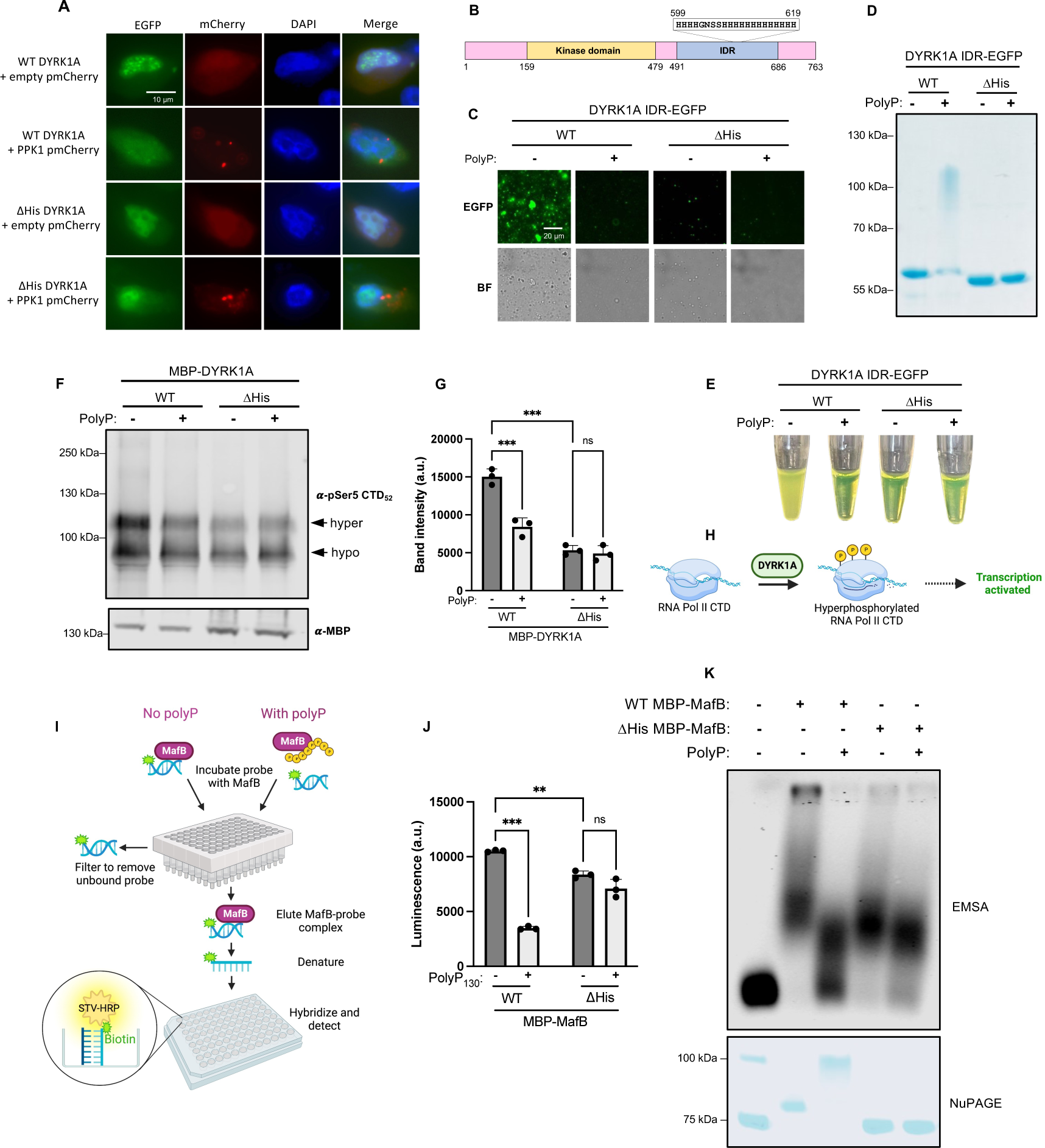
Polyphosphorylation negatively regulates DYRK1A and MafB activities *in vitro*. (A) PolyP overproduction in HeLa cells results in the loss of DYRK1A nuclear speckles. HeLa cells were co-transfected with WT or ΔHis (residues 599-619 deleted) EGFP-DYRK1A, plus either PPK1-mCherry or mCherry alone. (B) Schematic of DYRK1A. Phase separation of purified wild-type or ΔHis (residues 599-619 deleted) DYRK1A IDR-EGFP pre-treated with polyP_700_ as indicated imaged via microscopy (C) or in tubes (E). (D) NuPAGE of samples from panels E/D. (F and G) Kinase activity of MBP-DYRK1A on GST-CTD_52_ substrate detected via western blotting against phosphorylated CTD_52_ following SDS-PAGE. ImageJ densitometry of the hyperphosphorylated band is graphed for three independent experiments. Two-way ANVOA; ns, *P* > 0.05; ***, P < 0.001. (H) Schematic of DYRK1A activation of transcription. (I) Schematic of the Signosis filter plate method used for assessing MafB binding to its biotinylated DNA target. (J) Luminescent quantification of purified MBP-MafB binding to target DNA via the Signosis filter plate after being treated with polyP_130_. The ΔHis MafB in panels J and K lacks the histidine repeat region (residues 131-167). Two-way ANOVA; ns, *P* > 0.05; **, *P* ≤ 0.01, ***, P ≤ 0.001. (K) Agarose gel EMSA of MBP-MafB binding to IR700Dye-labelled DNA probe after being treated with polyP_130_.

The histidine repeat region of DYRK1A also drives phase separation *in vitro.*^36^ We purified recombinant DYRK1A intrinsically disordered region (IDR; residues 491-686) (Figure 4B), which contains the histidine repeat region, fused to EGFP. Upon addition of PEG 8000 as a crowding agent, WT DYRK1A IDR-EGFP formed phase-separated droplets visible both under the microscope and as turbidity in the reaction tubes. PolyP treatment of the WT protein reduced phase separation to a level comparable to the ΔHis sample (Figure 4C). Polyphosphorylation of the WT but not ΔHis protein was confirmed via NuPAGE gel (Figure 4D). A similar histidine-dependent disruption of phase separation by polyphosphorylation was observed for the cyclin T1 IDR (Figures S4C and S4D).

As a kinase, DYRK1A phosphorylates substrates at serine or threonine residues.^37^ One particularly important substrate of DYRK1A is the C-terminal domain (CTD) of the RPB1 subunit of human RNA polymerase II, which is phosphorylated within YSPTSPS repeats at Ser2 and Ser5.^29^ We used glutathione-S-transferase (GST)-CTD_52_ (RPB1 CTD containing all 52 heptapeptide repeats) as a substrate^36^ and detected phosphorylation with an anti-phospho-Rbp1 CTD Ser5 antibody (Figure S4E). DYRK1A phosphorylation generated two distinct states of phospho-CTD_52_: hyper- and hypo-phosphorylated, as previously described.^36^ The hyperphosphorylated state is critical for transcriptional elongation.^36^ PolyP treatment of WT MBP-DYRK1A reduced the amount of hyperphosphorylated CTD_52_ to levels on par with those of ΔHis MBP-DYRK1A. PolyP treatment did not significantly alter CTD_52_ hyperphosphorylation by ΔHis MBP-DYRK1A (Figures 4F–4H).

### iH-PPn inhibits MafB transcription factor activity *in vitro*

MafB is a transcription factor that binds DNA containing Maf-recognition elements (MAREs).^38^ We hypothesized that MafB attachment to another long negative biopolymer such as polyP may disrupt DNA binding. We tested this activity using both traditional gel electrophoretic mobility shift assay (EMSA) and a filter plate method whereby only DNA probe that is bound to MafB is retained and subsequently quantified via luminescent detection (Figure 4I). We confirmed MafB specificity for the labelled probe and ensured that polyP was not interfering with the filter plate assay (Figures S4F-S4G). In both the filter assay (Figure 4J) and gel EMSA (Figure 4K), polyP treatment of the WT MafB, but not the ΔHis, resulted in impaired DNA binding. The filter assay detected only a slight decrease in activity of the untreated ΔHis protein versus the untreated WT, indicating that the histidine repeat of MafB is not critical for DNA binding in the absence of polyP. Importantly, polyP treatment of the ΔHis protein yielded no significant additive inhibition of DNA binding, implicating the histidine repeat as the mediator of this phenomenon.

## DISCUSSION

The complexity of higher organisms is greatly enhanced by protein post-translational modification (PTM). Lysine polyphosphorylation (K-PPn) is a recently discovered PTM wherein polyP is attached to lysine residues via what is posited to be a covalent P-N bond.^12,13^ PolyP is attached so robustly during K-PPn that proteins exhibit a marked mobility shift upon NuPAGE electrophoresis, which is strongly denaturing.

Proteins that interact non-covalently with polyP would not be expected to shift in such harsh conditions, and indeed, NuPAGE has previously been used to differentiate between covalent and non-covalent polyP binding.^15^ We present a new interaction between polyP and proteins containing histidine repeats that blurs the NuPAGE-based distinction between covalency and non-covalency. We discovered 30 human and yeast proteins containing between 5-13 consecutive histidine residues that exhibited a polyP-dependent NuPAGE shift. Histidine deletion abolished the shift, and histidine-only peptide was sufficient to generate a shift, providing strong evidence that histidine residues are the direct target of polyP in this modification.

The absence of a phoshoramidate bond in the ^31^P NMR of polyphosphorylated 13H peptide, along with the sensitivity of polyP-polyhistidine binding towards pH and ionic strength strongly argue that this modification is ionic rather than covalent. The pH of 7.5 above which the polyP interaction is abolished coincides with the p*K*_a_ threshold at which histidine sidechains lose all positive charge. The sidechain p*K*_a_ of monomeric histidine is approximately 6, though this can vary considerably depending on residue environment.^39^ To describe the unique properties of this interaction, we propose the term ionic histidine polyphosphorylation (iH-PPn).

While positive charge therefore appears to be important for iH-PPn, it is clearly not the only factor involved, since replacement of the 13-histidine tract in Snf1 with 13 arginine residues was unable to rescue the shift. The considerably higher sidechain p*K*_a_s of arginine and lysine (12.5 and 10.5, respectively) would presumably provide even stronger electrostatic interaction with polyP, but the unique sidechain geometry of histidine may make it more suitable for polyP binding. Further studies are required to structurally characterize the histidine-polyP interaction interface.

Though none of our histidine repeat hits appear to overlap with hits from previous polyP interactome screening studies,^15,40,41^ our findings may warrant a closer look at the sequences of these proteins. For instance, the Disabled-1 (DAB1) protein discovered to interact with polyP in a recent screen^41^ contains 3 consecutive histidine residues (HHHAVH). It is therefore tempting to speculate that this histidine repeat may be involved in DAB1 polyP binding.

Our functional testing of DYRK1A and MafB following polyphosphorylation suggest that this modification may have widespread regulatory roles. Polyphosphorylation inhibited the *in vitro* phase separation, nuclear speckle formation, and kinase activity of DYRK1A in a histidine repeat-dependent manner, phenocopying the defects observed in ΔHis DYRK1A^23,36^. Traditional covalent PTMs such as methylation^42^ or phosphorylation^43^ within the low-complexity regions that drive condensate formation are well-established disruptors of phase separation. Non-covalent iH-PPn may disrupt DYRK1A phase separation and activity in a similar manner to these PTMs, either by steric occlusion of the low-complexity (histidine) region or alteration of the protein structure. The transition of DYRK1A from organized nuclear speckles to a diffuse nuclear/cytoplasmic localization upon polyP overproduction mirrors the changes in Nsr1 and Top1 localization upon K-PPn. Without polyphosphorylation, Nsr1 and Top1 are confined to the nucleolus, while K-PPn results in diffuse nuclear localization of both proteins.^12^

In addition to disrupting the functions of histidine repeats proper, we also show that iH-PPn can compromise protein function that is otherwise independent of the histidine repeat. The consecutive histidine tract of MafB is dispensable for its DNA binding activity and is relatively distant from its leucine zipper DNA binding motif. However, a polyP chain hundreds of phosphate units in length tethered to the histidine repeat could ostensibly wrap around the protein to competitively block DNA binding or otherwise disrupt protein structure. This scenario is reminiscent of the inhibition of Top1 DNA-relaxing topoisomerase activity by K-PPn.^12^ The 8-histidine repeat (H131-H138) of MafB is highly conserved in vertebrates,^44^ and a missense mutation of the first residue in this repeat (H131Q) has been associated with orofacial clefts.^44,45^

While we focused on proteins with long (≥ 5 consecutive) histidine stretches, the example of HRG suggests that iH-PPn may also occur on proteins with shorter interrupted histidine-rich regions, which could vastly expand the scope of target proteins. Our observation that iH-PPn can occur on 6-His tracts and interfere with binding to Ni^2+^ resin also serves as a cautionary note to scientists using recombinant His-tagged proteins for polyP research or biotechnological applications where polyP may be present. The pH- and salt-sensitivity of this modification raises the intriguing possibility for spatiotemporal fine-tuning of polyP attachment. Cells are capable of exquisitely sensitive control of pH and ionic strength, which we envision could serve as an additional layer of regulation to control if, when, or where proteins undergo iH-PPn. The extent to which iH-PPn occurs *in vivo*, and what cellular factors regulate this phenomenon, remain open questions for future research. Our work provides a foundation for studying histidine-polyP interactions and their potential regulatory roles in greater detail.

### Limitations of the Study

The promiscuous functions of polyP make it difficult to ascertain that some of the effects we observed are directly due to iH-PPn. PolyP possesses non-specific chaperone activity,^5^ the influence of which we cannot exclude in the case of phase separation. PPK1 overexpression in HeLa cells is known to cause pleiotropic abnormalities,^46^ which may have contributed to the loss of DYRK1A nuclear speckles. Since these phenotypes are already abolished for ΔHis proteins, we could not rely on classical epistasis experiments (i.e., a lack of response to polyP treatment of ΔHis mutants) to confirm a causal relationship. It also remains to be determined if the quantities of endogenous polyP in human cells are sufficient to cause iH-PPn, and whether this interaction occurs naturally *in vivo*.

## STAR METHODS

### Plasmids and mutagenesis

A detailed description of the plasmids used in this study is available in Table S2. Insertion and deletion mutagenesis was performed via PCR as described.^47^ Successful mutation was verified by Plasmidsaurus whole-plasmid sequencing.

### Bacterial protein expression and purification

Proteins were expressed from BL21(DE3) RIPL *E. coli* cells (Agilent). Overnight cultures were inoculated into fresh LB at 1/100 dilution and grown at 37 °C until OD_600_ = 0.6, at which point expression was induced with 1 mM IPTG. Cultures were grown at 20 °C overnight. Cells were pelleted and resuspended in lysis buffer comprised of 50 mM Tris pH 8, 250 mM NaCl, 5% glycerol, 3 mM 2-mercaptoethanol. Cells were lysed via sonication with a Branson instrument for 6 minutes, with cycles of 5s on and 15s off. Lysates were clarified via centrifugation at 18,000 rpm for 30 min. Clarified supernatant was applied to the appropriate resin (amylose for MBP tags, NEB E8021L; glutathione for GST tags, G-Biosciences 786-310; nickel-NTA for His-tags, Cytiva 17057502). After washing with lysis buffer, proteins were eluted with either 10 mM maltose, 10 mM reduced glutathione, or 200 mM imidazole depending on tag.

### In vitro polyphosphorylation assays

Purified proteins or cell lysates were incubated with polyP for 30 min at RT. Unless otherwise indicated, the polyP and the negative control used throughout this paper were polyP_700_ (Kerafast EUI003) and monomeric NaH_2_PO_4_, respectively, both added to 5 mM final concentration. All polyP concentrations are in terms of P_i_ monomers. Samples were mixed 1:1 with 2x Leammli sample buffer (120 mM Tris pH 6.8, 100 mM DTT, 20% glycerol, 4% SDS, 0.2% bromophenol blue) and boiled at 95°C for 5 minutes.

Samples were run on 4-12% Bis-Tris NuPAGE gels (ThermoFisher NP0336BOX) at 200 V for 45 minutes in 1X NuPAGE running buffer (ThermoFisher NP0001) or a homemade equivalent composed of 50 mM Tris, 50 mM MOPS, 1 mM EDTA, 5 mM sodium bisulfite, and 0.1% SDS.^13^ For pulldown (Figure 2C), MBP-Snf1(1-65) was immobilized on amylose beads at the indicated pH, treated with 5 mM polyP_700_, washed 4x with buffer of indicated pH, then eluted by boiling in Laemmli buffer.

For NuPAGE shift test in the presence of other negatively charged polymers, polyanion concentrations are in terms of monomer constituents adjusted based on the average charge per formula unit. The formula weights used for these calculations are as follows: polyP, 102 g/mol; dextran sulfate trisodium salt (Sigma D-8906), 468 g/mol; heparin (Millipore 375095) (considering a disaccharide of uronic acid and glucosamine as a monomer) 593.45 g/mol; single-stranded DNA (Millipore D7656), 325 g/mol. Concentrations were then normalized to an average charge of −1 per formula unit. For example, polyP possesses −1 charge per monomer, so it required no normalization. Dextran sulfate possesses −3 charge per monomer, thus it was diluted by 1/3.

### Western blotting

Proteins were transferred to Immobilon-FL PVDF membranes (Millipore IPFL00010) for immunoblotting. The primary antibodies used in this study are as follows: mouse anti-MBP (Cell Signalling 2396S), rabbit anti-GFP (Cell Signalling 2956S), rabbit anti-P-Rbp1 CTD Ser5 (Cell Signalling 13523S), rabbit anti-cyclin T1 (GeneTex GTX133413), and rabbit anti-YY1 (GeneTex GTX110625). Unless otherwise specified the secondary antibodies used were goat anti-mouse DyLight 680 (Invitrogen 35518) and goat anti-rabbit DyLight 800 (Invitrogen SA535571). Blots were imaged using a LI-COR Odyssey CL-x.

For Figure S1D only, the primary antibody was mouse anti-GFP (Living Colors 632381) and the secondary antibody was goat anti-mouse HRP conjugate (BioRad 172-1011). Blots were developed with Immobilon ECL reagent (Sigma WBKLS0500) and autoradiography film from Hyblot CL (8X10).

### Peptides and NMR

Peptides were synthesized by Royobiotech (Shanghai). NMR samples were prepared with 0.5 mM untagged (no FITC) peptide dissolved in 50 mM Tris pH 7.5, 10% D2O, and mixed with 5 mM polyP_700_ for 1 h at room temperature. The pHis positive control was chemically synthesized from poly-L-histidine (Sigma P9386), triethylamine (Sigma 471283), and phosphoryl chloride (Sigma 8223391000) as described,^33^ then dialyzed into 50 mM Tris pH 7.5, 10% D2O. Spectra were obtained using a Bruker Neo 700-MHz spectrometer operating at 283.52 MHz for ^31^P.

### Circular dichroism

For CD, FITC-tagged peptides were dissolved in ultrapure water at 0.25 mM. Where indicated, polyP_700_ was added to 2.5 mM and incubated for 30 min at RT. Samples were run on a Chirascan (AppliedPhotophysics) CD spectrometer. Samples were scanned in a 0.1 mm pathlength cuvette between 180 and 260 nm at RT.

### Size exclusion chromatography

To monitor polyP binding in solution, samples were fractionated by FPLC using a Superdex 200 Increase 10/300 GL (Cytiva 28990944) size exclusion column. For initial experiments, the column was equilibrated and run with buffer composed of 50 mM Tris, 50 mM MOPS, 1 mM EDTA, and 5 mM sodium bisulfite, final pH 7.5 to mimic NuPAGE gel running buffer. For salt disruption experiments, the indicated concentrations of NaCl were added to the above running buffer recipe. For pH experiments, pH was adjusted by varying the ratio of Tris to MOPS, thus avoiding concentrated HCl or NaOH which could confound ionic strength. In addition to 1 mM EDTA and 5 mM sodium bisulfite, the various pH-varied buffers contained Tris/MOPS ratios as follows: pH 6.2, 5.6 mM Tris, 50 mM MOPS; pH 7.75, 50 mM Tris, 42.1 mM MOPS; pH 8, 50 mM Tris, 30.6 mM MOPS; pH 9.15, 50 mM Tis, 0 mM MOPS.

### HeLa cell transfection, imaging, and protein extraction

HeLa cells were cultured at 37°C with 5% CO_2_ in Dulbecco’s Modified Eagle Medium (DMEM) with 10% fetal bovine serum. Cells were seeded onto glass coverslips pretreated with fibronectin in 24-well plates and transfected at ∼ 80% confluency using Lipofectamine 3000 (ThermoFisher) as per the manufacturer instructions. Cells were harvested 48 h post transfection, fixed with 4% paraformaldehyde in PBS for 10 min at RT, permeabilized with 0.1% Triton X-100, with three 5 min washed with PBS between each step. Coverslips were mounted with ProLong Diamond containing DAPI (ThermoFisher P36962). Protein localization was observed via an Olympus IX83 microscope with 100x oil objective.

To harvest protein extracts, cells were washed 3x in PBS then scraped into TNTE-FB lysis buffer (20 mM Tris pH 7.5, 150 mM NaCl, 5 mM EDTA, 0.3% Triton X-100, 10 mM NaF, 40 mM β-glycerophosphate) supplemented with Halt protease inhibitor cocktail (ThermoFisher 78430). Cells were incubated on ice for 30 min to allow lysis, then centrifuged 3 min at 10,000 rpm to pellet debris. Supernatant was mixed 1:1 with 2x Laemmli loading buffer and boiled for 5 min prior to gel electrophoresis.

### In vitro phase separation

*In vitro* phase separation experiments of DYRK1A IDR-EGFP and Cyclin T1 IDR-EGFP were performed as described,^36^ with slight modification. Briefly, purified proteins were exchanged into buffer containing 150 mM NaCl, 20 mM Tris-HCl pH 7.5, and 1 mM DTT via size exclusion and concentrated to 20 mg/mL. To facilitate the formation of phase-separated liquid droplets, the fusion proteins were diluted into buffer containing 20 mM Tris-HCl pH 7.5, 1 mM DTT, and 37.5 mM NaCl. The DYRK1A IDR-EGFP fusions were diluted to 2.7 mg/mL final concertation, and the cyclin T1 IDR-EGFP fusions were diluted to 6 mg/mL final concentration. The DYRK1A solutions were supplemented with 10% PEG8000 final concentration. 5 µL of solution was trapped between coverslips and imaged on an Olympus IX83 epifluorescence microscope.

### DYRK1A protein kinase assay

The DYRK1A protein kinase activity assay was adapted from published protocols ^29^. The GST-CTD_52_ substrate was derived from Lu et al. and consisted of residues 1589 to 1970 of human RNA polymerase II RPB1 subunit (Uniprot P24928) fused at its N-terminus to a GST tag ^36^. MBP-DYRK1A bound to amylose beads was incubated with 5 mM polyP_700_ for 30 min at RT, then washed 3x with buffer containing 50 mM MOPS, 50 mM Tris, 1 mM EDTA and 5 mM sodium bisulfite, pH 7.5. Washed proteins were eluted from beads in phosphorylation buffer (50 mM HEPES pH 7.4, 10 mM MgCl_2_, 10 mM MnCl_2_, 1 mM DTT, and 100 µM ATP) supplemented with 10 mM maltose. Purified GST-CTD_52_ was also exchanged into phosphorylation buffer. Approximately 30 ng of MBP-DYRK1A in phosphorylation buffer was mixed with 200 ng of GST-CTD_52_ in phosphorylation buffer for 30 min at 30 °C. Reactions were quenched by boiling with Laemmli loading buffer for 5 min, then subjected to SDS-PAGE followed by immunoblotting with anti-pSer5 antibody to detect GST-CTD_52_ phosphorylation.

### TF filter plate and EMSA DNA binding assays

For both plate-based and gel-based transcription factor DNA binding assays, purified MBP fusion proteins in NuPAGE-mimic buffer (50 mM Tris, 50 mM MOPS, 1 mM EDTA, and 5 mM sodium bisulfite, final pH 7.5) were incubated with 5 mM polyP_130_ for 30 min at RT. We used polyP_130_ for these assays since it more closely mimics the short chain polyP observed in mammalian platelets,^48^ and because it was easier to separate unbound polyP_130_ from our proteins via SEC compared to the longer polyP_700_. Proteins were then subjected to SEC on a Superdex S200 16/600 column equilibrated in NuPAGE-mimic buffer to remove any unbound polyP. Protein-containing fractions were pooled, and concentrations were normalized via Bradford assay. The TF filter plate assay kit was purchased from Signosis (FA-0010) and contained the custom Maf recognition element (MARE) probe 5′-AGCTCGGAATTGCTGACTCATCATTACTC-3′ derived from previous MafB studies.^49,50^ Approximately 0.25 µg of protein was added per reaction well, and assay was conducted as prescribed by Signosis.

For gel-based EMSA, proteins were prepared as above, and probe labelled with IRDye® 700 was used to visualize binding. Labelled forward probe was based on the MafB MARE with sequence 5’-IRDye® 700-TGCTGACTCAGCA-3’, which was annealed to unlabelled reverse sequence 5’-TGCTGAGTCAGCA-3’ to create double-stranded probe. Binding reaction recipe was adapted from LI-COR (https://licor.app.boxenterprise.net/s/1mqrtw8bbwd31jgcpk5r); 20 µL reactions contained 2 µL of 10X binding buffer (100 mMTris, 500 mM KCl, 10 mM DTT; pH 7.5), 2 µL of 25 mM DTT/2.5%Tween 20, 1 µL of probe (0.2 µM stock), 1 µL of 100 mM MgCl_2_, 8 µL protein (yielding approximately 0.5 µg per reaction), and 6 µL water. Reactions were incubated 30 min at RT, then mixed with 4 µL of 6X orange loading dye. Samples were electrophoresed on 1% agarose gel in 0.5X TBE buffer, pH 7.0. Gels were imaged on a LI-COR Odyssey CL-x using the 700 nm channel.

### Statistical analyses

All results are expressed as the mean *±* standard deviation of three independent experiments unless otherwise stated. T-test and two-way ANOVA were performed in GraphPad Prism V9 to determine statistical significance.

## ACKNOWLEDGEMENTS

We thank Dr. Susana De la Luna (Centre for Genomic Regulation, Barcelona) for providing the pEGFP DYRK1A construct and Dr. Andrew Craig (Queen’s University) for providing pmCherry. PolyP standards (P14, P60, and P130) were a generous gift from Dr. Toshikazu Shiba (RegeneTiss). We also thank Dr. Hongwei Tan (Beijing Normal University) for help with pKa calculations, and Dr. Françoise Sauriol (Queen’s University) for her NMR assistance. Cartoon figures were created using BioRender.

## AUTHOR CONTRIBUTIONS

Conceptualization, Z.J., N.N., and K.L.; Methodology, N.N., K.L., Z.Y., and M.D.; Investigation, N.N., K.L, K.K, A.D., and M.D.; Writing – Original Draft, N.N.; Writing – Review and Editing, N.N., Z.J., K.L., and M.D.; Funding Acquisition, Z.J.; Supervision; Z.J.

