## Supplemental Figures for "Ionic polyphosphorylation of histidine repeat proteins by inorganic polyphosphate"

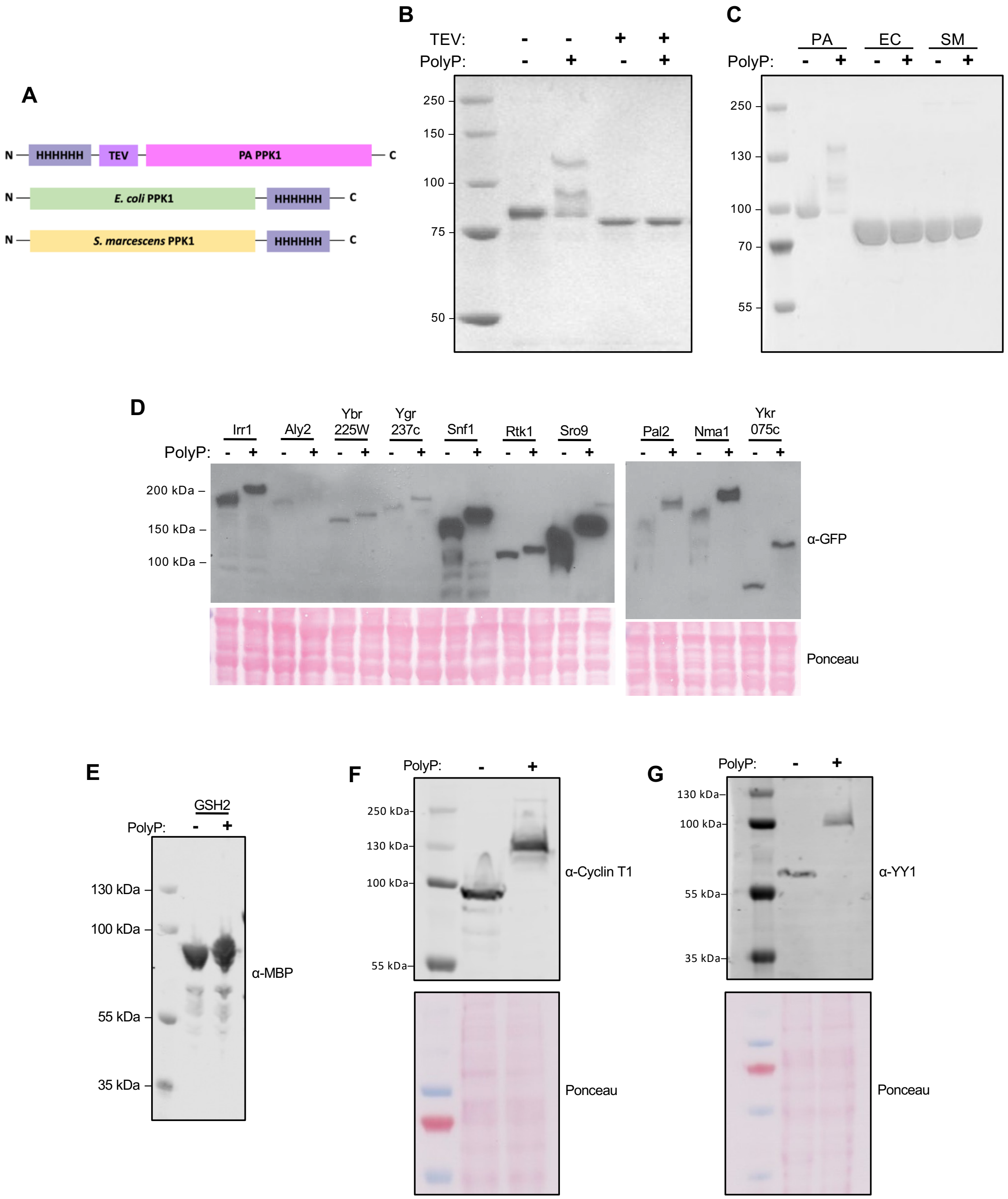

**Figure S1. Histidine repeats confer a NuPAGE shift**. Related to Figure 1. (A-C) Hexahistidine tag confers a polyP-dependent NuPAGE shift to *Pseudomonas aeruginosa* (PA) PPK1, but not *Escherichia coli* (EC) or *Serratia marcescens* (SM) PPK1. (A) Schematic of PPK1 constructs tested. TEV represents the cleavage site for tobacco etch virus (TEV) protease. (B) PA PPK1 was treated with TEV protease, followed by 5 mM polyP$_{700}$ as indicated. (C) PPK1 from different bacterial species were treated with 5 mM polyP$_{700}$ as indicated. Proteins were resolved on NuPAGE gels and stained with Coomassie blue. (D) Screening *Saccharomyces cerevisiae*

histidine repeat proteins for NuPAGE shift. Lysates were prepared from a GFP-tagged library in which each yeast open reading frame is expressed from its genomic locus as a GFP fusion. Lysates were treated with either 10 mM polyP$_{700}$ or water, resolved via NuPAGE, transferred to PVDF, and immunoblotted with anti-GFP antibody. (E) NuPAGE shift test of recombinant MBP-tagged human GSH2 (overflow from Figure 1B) visualized via anti-MBP immunoblot in *E. coli* expression lysate treated with or without polyP$_{700}$. (F and G) Immunoblots against endogenous untagged cyclin T1 and YY1 in HeLa lysates treated with or without 5 mM polyP$_{700}$ and resolved on NuPAGE.

**Table S1**. *S. cerevisiae* GFP-tagged library His repeat protein NuPAGE screening summary

| Name | Uniprot ID | Activity | Maximum # consecutive His residues | MW alone (kDa) | MW with GFP (kDa) | Shift? |
|---|---|---|---|---|---|---|
| **Irr1** | P40541 | Component of cohesin complex | 2 | 133 | 160 | Yes |
| **Aly2** | P47029 | Alpha arrestin, Ub-ligase adaptor | 2 | 117 | 139 | Yes |
| **Ybr225W** | P38321 | Unknown | 4 | 101 | 128 | Yes |
| **Ygr237c** | P50089 | Unknown | 7 | 89 | 116 | Yes |
| **Snf1** | P06782 | AMP-activated S/T protein kinase | 13 | 72 | 99 | Yes |
| **Rtk1** | Q12100 | Putative S/T protein kinase | 7 | 70 | 97 | Yes |
| **Sro9** | P25567 | RNA-binding protein | 3 | 48 | 75 | Yes |
| **Pal2** | P38809 | Involved in clathrin-mediated endocytosis | 4 | 41 | 68 | Yes |
| **Nma1** | Q06178 | Adenylyltransferase | 7 | 46 | 73 | Yes |
| **Ykr075c** | O60481 | Unknown | 6 | 36 | 63 | Yes |
| **Num1** | Q00402 | Nuclear migration | 2 | 312 | 339 | No |
| **Pmd1** | P32634 | Putative negative regulator of early meiotic gene | 6 | 195 | 222 | No |
| **Mhp1** | P43638 | Microtubule-associated protein | 5 | 155 | 182 | No |
| **Rom2** | P51862 | Guanine nucleotide exchange factor | 8 | 153 | 180 | No |
| **Duf1** | Q99247 | Ubiquitin-binding protein | 9 | 125 | 152 | No |
| **Pbp1** | P53297 | Glucose deprivation induced stress granules | 2 | 79 | 106 | No |
| **Dot6** | P40059 | rRNA and ribosome biogenesis | 3 | 73 | 100 | No |
| **Gyp1** | Q08484 | Cis-golgi GTPase-activating protein | 4 | 73 | 100 | No |
| **Vts1** | Q08831 | DNA- and RNA-binding protein | 4 | 56 | 83 | No |
| **Hsp42** | Q12329 | Heat shock protein | 2 | 43 | 70 | No |
| **Opy1** | P38271 | Unknown | 7 | 38 | 65 | No |
| **Tph3** | P47072 | Unknown | 2 | 63 | 90 | No |
| **Fps1** | P23900 | Aquaglyceroporin, plasma membrane channel | 4 | 74 | 101 | No |
| **Erg25** | P53045 | C-4 methyl sterol oxidase | 3 | 36 | 63 | No |

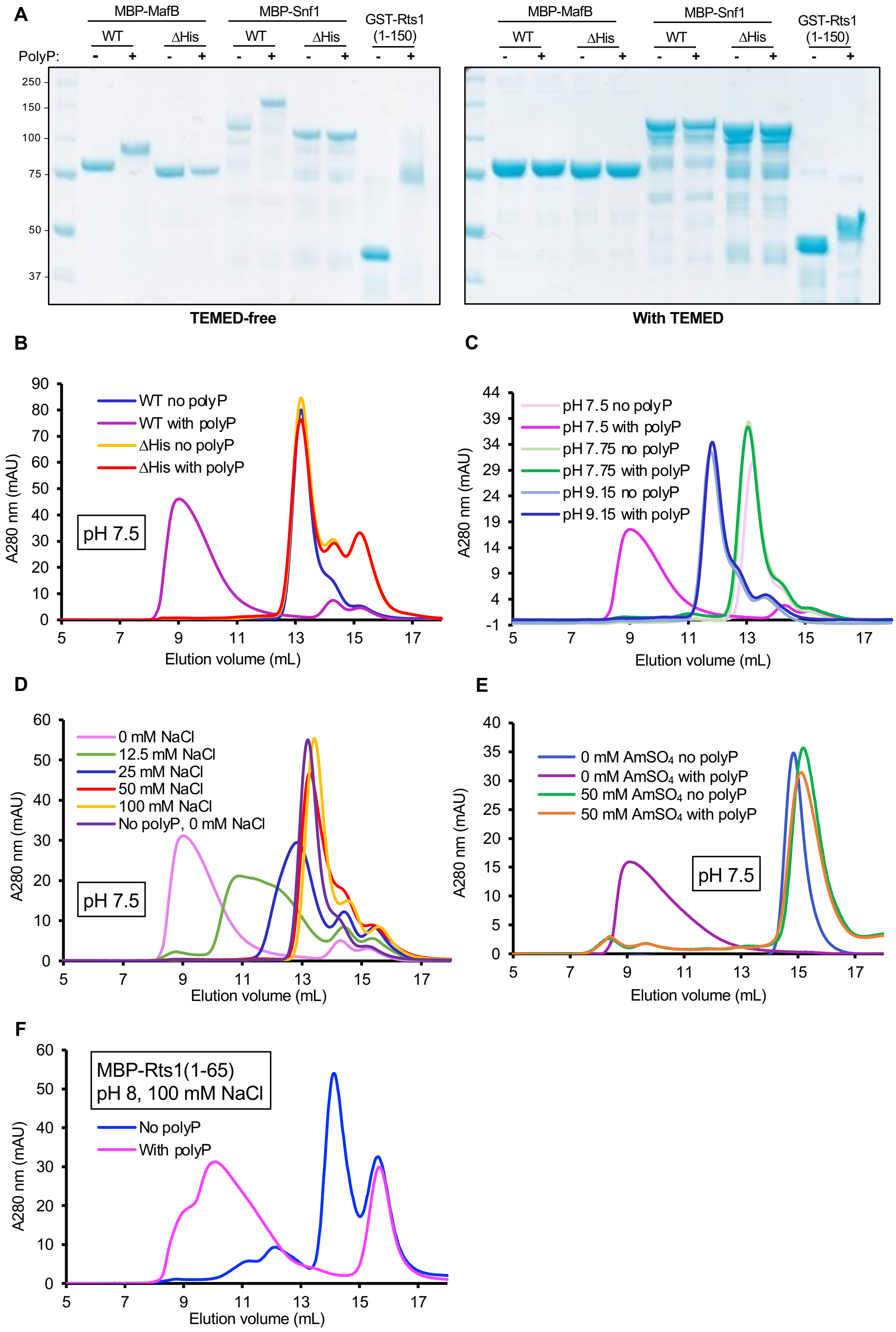
A
MBP-MafB
WT
ΔHis
MBP-Snf1
WT
ΔHis
GST-Rts1 (1-150)
PolyP:
250
150
100
75
50
37
TEMED-free
With TEMED
B
WT no polyP
WT with polyP
ΔHis no polyP
ΔHis with polyP
pH 7.5
A280 nm (mAU)
Elution volume (mL)
C
pH 7.5 no polyP
pH 7.5 with polyP
pH 7.75 no polyP
pH 7.75 with polyP
pH 9.15 no polyP
pH 9.15 with polyP
A280 nm (mAU)
Elution volume (mL)
D
0 mM NaCl
12.5 mM NaCl
25 mM NaCl
50 mM NaCl
100 mM NaCl
No polyP, 0 mM NaCl
pH 7.5
A280 nm (mAU)
Elution volume (mL)
E
0 mM AmSO4 no polyP
0 mM AmSO4 with polyP
50 mM AmSO4 no polyP
50 mM AmSO4 with polyP
pH 7.5
A280 nm (mAU)
Elution volume (mL)
F
MBP-Rts1(1-65)
pH 8, 100 mM NaCl
No polyP
With polyP
A280 nm (mAU)
Elution volume (mL)

**Figure S2. Sensitivity of histidine polyphosphorylation to TEMED, pH, and ionic strength**. Relates to Figure 2. (A) TEMED collapse of histidine polyphosphorylation. Homemade Bis-Tris gels were polymerized and run with identical samples under identical conditions except for the addition of tetramethylethylenediamine (TEMED) at 1 μL/mL. GST-Rts1(1-150) included as a PASK-containing K-PPn positive control. (B-D) Size exclusion of MBP-MafB(80-232) in the presence or absence of 5 mM polyP$_{700}$. (B) NuPAGE buffer containing 0 mM NaCl at pH 7.5. ΔHis MBP-MafB(80-232) is deleted in both histidine repeat regions (residues 131-167 deleted). (C) WT MBP-MafB(80-232) in NuPAGE buffer containing 0 mM NaCl at varied pH. (D) WT MBP-MafB(80-232) in NuPAGE buffer at pH 7.5 with varied [NaCl]. (E) MBP-Snf1(1-65) was incubated with or without polyP as above and subjected to SEC in buffer containing no salt (0 mM AmSO$_4$) or buffer supplemented with 50 mM ammonium sulfate (50 mM AmSO$_4$). (F) The PASK-containing positive control MBP-Rts1(1-65) exhibits a size exclusion shift in NuPAGE buffer containing 150 mM NaCl, pH 8. Proteins incubated with 5 mM polyP$_{700}$ where indicated for all the above SEC experiments.

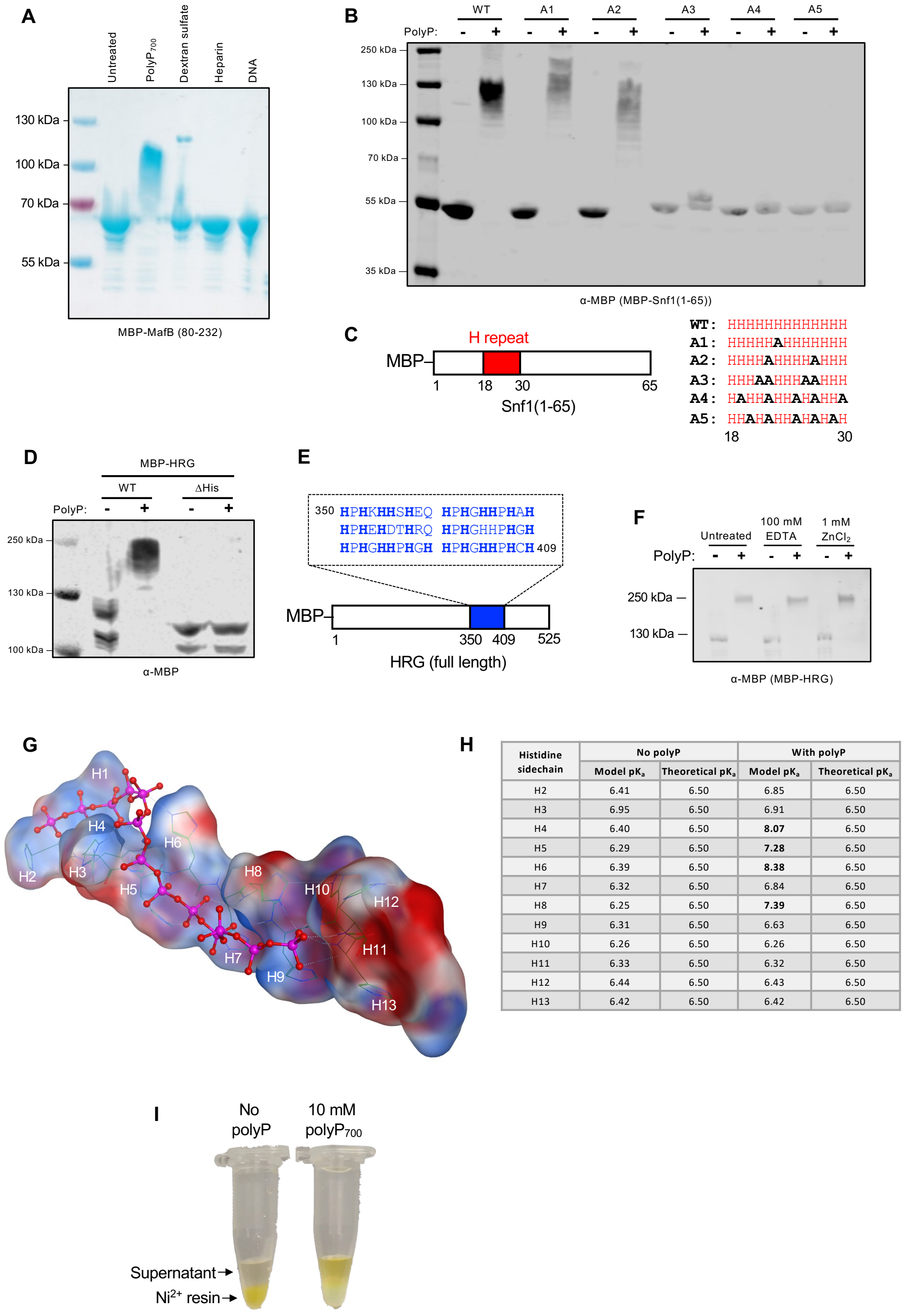

| Histidine sidechain | No polyP | | With polyP | |
|---|---|---|---|---|
| | Model $pK_a$ | Theoretical $pK_a$ | Model $pK_a$ | Theoretical $pK_a$ |
| H2 | 6.41 | 6.50 | 6.85 | 6.50 |
| H3 | 6.95 | 6.50 | 6.91 | 6.50 |
| H4 | 6.40 | 6.50 | **8.07** | 6.50 |
| H5 | 6.29 | 6.50 | **7.28** | 6.50 |
| H6 | 6.39 | 6.50 | **8.38** | 6.50 |
| H7 | 6.32 | 6.50 | 6.84 | 6.50 |
| H8 | 6.25 | 6.50 | **7.39** | 6.50 |
| H9 | 6.31 | 6.50 | 6.63 | 6.50 |
| H10 | 6.26 | 6.50 | 6.26 | 6.50 |
| H11 | 6.33 | 6.50 | 6.32 | 6.50 |
| H12 | 6.44 | 6.50 | 6.43 | 6.50 |
| H13 | 6.42 | 6.50 | 6.42 | 6.50 |

**Figure S3. Specificity, *in silico* modelling, and nickel purification interference of iH-PPn.** Relates to Figures 2 and 3. (A) Other biological polyanions are unable to replicate the marked polyP-induced NuPAGE shift of histidine-repeat protein MBP-MafB(80-232). All polymers added to 3.75 mM final concentration (in terms of monomers adjusted such that each monomeric unit has -1 charge). (B) NuPAGE shift test of MBP-Snf1(1-65) following alanine replacement within the 13H repeat as shown in panel (C). (D and E) MBP-HRG exhibits a NuPAGE shift that is abolished upon deletion of the histidine-rich region (residues 350-409). (F) The MBP-HRG shift is unaffected by pre-treatment with large molar excess of EDTA or zinc. (G) Docking model of 13-His peptide in complex with $polyP_{13}$ generated in Molecular Operating Environment (MOE). Energy minimized 13-His and $polyP_{13}$ were set as the receptor and ligand, respectively. (H) Predicted $pK_a$ values of the histidine sidechains in the presence or absence of polyP were calculated using PROPKA software for the docked model and tabulated. (I) Pre-treatment of FITC-13H peptide prevents its binding to $Ni^{2+}$ resin.

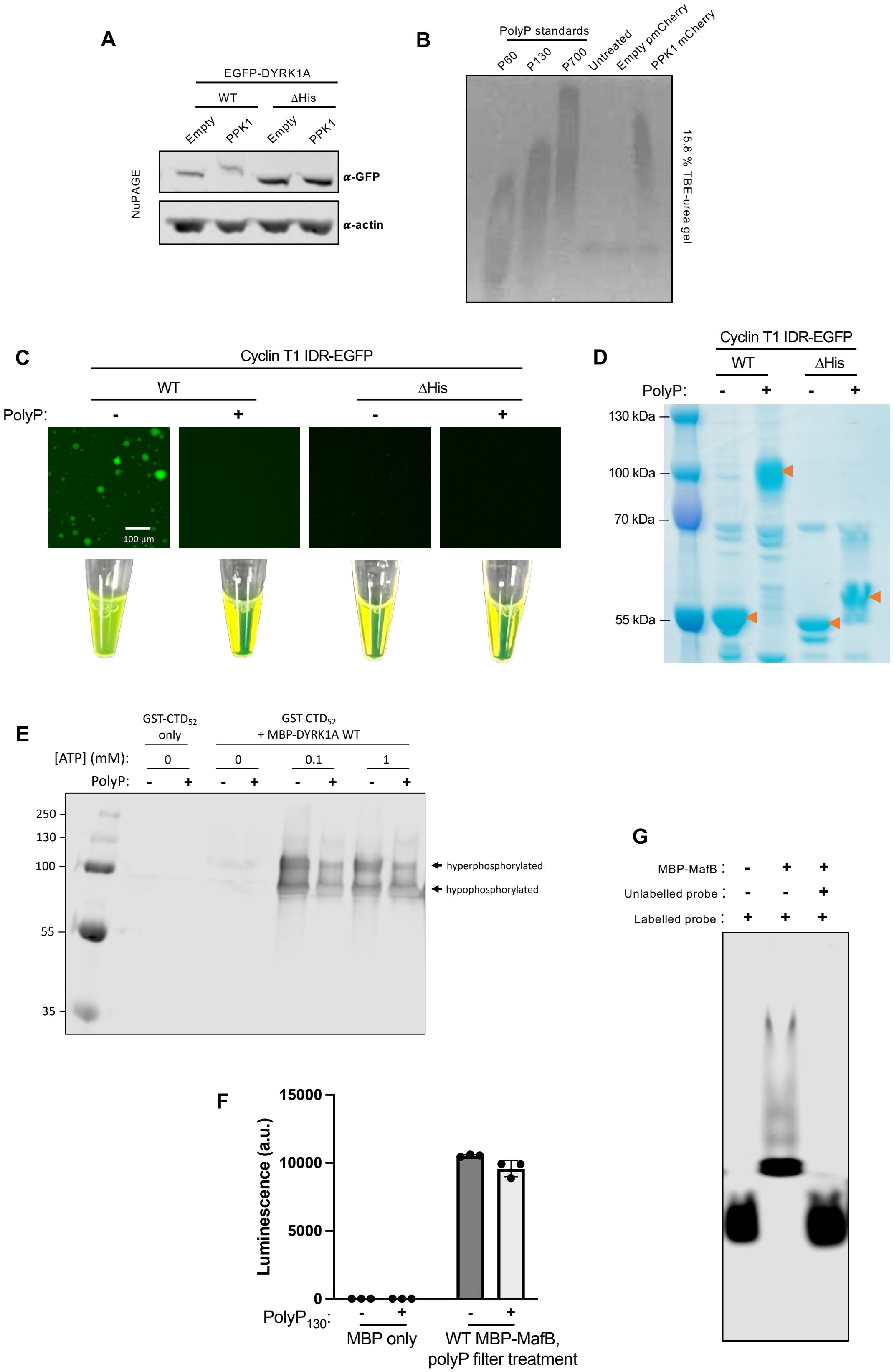
A
EGFP-DYRK1A
WT
ΔHis
Empty
PPK1
Empty
PPK1
NuPAGE
α-GFP
α-actin
B
PolyP standards
P60
P130
P700
Untreated
Empty pmCherry
PPK1 mCherry
15.8 % TBE-urea gel
C
Cyclin T1 IDR-EGFP
WT
ΔHis
PolyP:
-
+
-
+
100 µm
D
Cyclin T1 IDR-EGFP
WT
ΔHis
PolyP:
-
+
-
+
130 kDa
100 kDa
70 kDa
55 kDa
E
GST-CTD52 only
GST-CTD52 + MBP-DYRK1A WT
[ATP] (mM):
0
0
0.1
1
PolyP:
250
130
100
55
35
hyperphosphorylated
hypophosphorylated
F
Luminescence (a.u.)
15000
10000
5000
0
PolyP130:
MBP only
WT MBP-MafB, polyP filter treatment
G
MBP-MafB :
Unlabelled probe :
Labelled probe :

**Figure S4. iH-PPn alters the activities of DYRK1A, cyclin T1, and MafB.** Relates to Figure 4. (A) HeLa lysates from Figure 4A were subjected to NuPAGE and immunoblotting to demonstrate that PPK1-mCherry generates sufficient polyP to polyphosphorylate WT EGFP-DYRK1A. (B) HeLa lysates from Figure 4A were subjected to urea PAGE and negative DAPI staining to visualize polyP produced by PPK1-mCherry. (C) Phase separation of purified cyclin T1 IDR (residues 462-654; encompasses the histidine repeat) fused to EGFP. Cyclin T1 residues 517-528 deleted in the ΔHis protein. (D) NuPAGE of cyclin T1 IDR-EGFP samples from panel (C) tubes. Orange arrows denote bands of interest. (E) Validation of the GST-CTD$_{52}$ substrate and anti-pSer5 CTD antibody as a means of assessing DYRK1A kinase activity. (F) Purified MBP alone has no binding activity in the MafB filter plate assay and polyP does not interfere with the plate. To exclude the possibility that polyP was interfering with the filter plate itself (and not just disrupting probe binding), the filter plate was pre-treated with 50 µM polyP$_{130}$, washed as directed, then an identical sample of polyP-free WT MBP-MafB from Figure 4J was applied to both the untreated and pre-treated wells. (G) EMSA gel specificity of the IRDye-labelled probe for MafB confirmed via competition assay with unlabelled DNA probe duplex of identical sequence added at 100-fold molar excess to the reaction mix. Competition eliminated labelled DNA binding, indicating that the binding is specific.

**Table S2.** Plasmids used in this study

| Storage name | Insert gene | Tag | Backbone vector | Description | Source |
|---|---|---|---|---|---|
| NN138 | N/A | N-MBP-TEV-insert-C | pET16b | Empty vector for expressing MBP alone | Jia lab |
| NN236 | MafA | N-MBP-TEV-insert-C | pET28a | Human WT MafA (Uniprot Q8NHW3) | Codon optimized for *E. coli* by Sangon Biotech |
| NN145 | MafB | N-MBP-insert-C | pET16b | Human WT MafB (Uniprot Q9Y5Q3) | |
| NN280 | MafB ΔHis | N-MBP-insert-C | pET16b | As above, but with residues 131-167 deleted | |
| NN228 | YY1 | N-MBP-TEV-insert-C | pET28a | Human WT YY1 (Uniprot P25490) | |
| NN230 | NLK | N-MBP-TEV-insert-C | pET28a | Human WT NLK (Uniprot Q9UBE8) | |
| NN227 | Cyclin T1 | N-MBP-TEV-insert-C | pET28a | Human WT cyclin T1 (Uniprot O60563) | |
| NN239 | DYRK1A | N-MBP-TEV-insert-C | pET28a | Human WT DYRK1A (Uniprot Q13627) | |

| | | | | |
|---|---|---|---|---|
| NN262 | DYRK1A ΔHis | N-MBP-TEV-insert-C | pET28a | As above, but with residues 599-619 deleted |
| NN240 | HAND1 | N-MBP-TEV-insert-C | pET28a | Human WT HAND1 (Uniprot O96004) |
| NN242 | PRIC3 | N-MBP-TEV-insert-C | pET28a | Human WT PRIC3 (Uniprot O43900) |
| NN243 | DLX2 | N-MBP-TEV-insert-C | pET28a | Human WT DLX2 (Uniprot Q07687) |
| NN244 | ZIC3 | N-MBP-TEV-insert-C | pET28a | Human WT ZIC3 (Uniprot O60481) |
| NN245 | NKD2 | N-MBP-TEV-insert-C | pET28a | Human WT NKD2 (Uniprot Q969F2) |
| NN246 | LRCH1 | N-MBP-TEV-insert-C | pET28a | Human WT LRCH1 (Uniprot Q9Y2L9) |
| NN247 | PLK2 | N-MBP-TEV-insert-C | pET28a | Human WT PLK2 (Uniprot Q9NYY3) |
| NN248 | VGLL3 | N-MBP-TEV-insert-C | pET28a | Human WT VGLL3 (Uniprot A8MV65) |
| NN249 | NFP2 | N-MBP-TEV-insert-C | pET28a | Human WT NFP2 (Uniprot Q7Z417) |
| NN250 | ANK57 | N-MBP-TEV-insert-C | pET28a | Human WT ANK57 (Q53LP3) |
| NN251 | MEPC | N-MBP-TEV-insert-C | pET28a | Human WT MEPC (Uniprot Q7L2J0) |
| NN252 | RHOB2 | N-MBP-TEV-insert-C | pET28a | Human WT RHOB2 (Uniprot Q9BYZ6) |
| NN253 | GATA6 | N-MBP-TEV-insert-C | pET28a | Human WT GATA6 (Uniprot Q92908) |
| NN254 | NR4A3 | N-MBP-TEV-insert-C | pET28a | Human WT NR4A3 (Uniprot Q92570) |
| NN234 | POU4F1 | N-MBP-TEV-insert-C | pET28a | Human WT POU4F1 (Uniprot Q01851) |
| NN229 | POU4F2 | N-MBP-TEV-insert-C | pET28a | Human WT POU4F2 (Uniprot Q12837) |
| NN232 | HOXA9 | N-MBP-TEV-insert-C | pET28a | Human WT HOXA9 (P31269) |
| NN233 | OTX1 | N-MBP-TEV-insert-C | pET28a | Human WT OTX1 (Uniprot P32242) |
| NN237 | MECP2 | N-MBP-TEV-insert-C | pET28a | Human WT MECP2 (Uniprot P51608) |
| NN238 | HRC | N-MBP-TEV-insert-C | pET28a | Human WT HRC (Uniprot P23327) |
| NN231 | GSH2 | N-MBP-TEV-insert-C | pET28a | Human WT GSH2 (Uniprot Q9BZM) |

| | | | | | |
|---|---|---|---|---|---|
| NN175 | HRG | N-MBP-TEV-insert-C | pET16b | Human WT HRG (Uniprot P04196) | Codon optimized for *E. coli* by GenScript |
| NN286 | HRG ΔHis | N-MBP-TEV-insert-C | pET16b | As above, but with residues 350-409 deleted | |
| NN261 | PolII $CTD_{52}$ | N-GST-PreScission-insert-C | pGEX6P1 | Residues 1589 to 1970 of human RPB1 (Uniprot P24928) | |
| NN208 | Rts1(1-150) | N-GST-PreScission-insert-C | pGEX6P1 | *S. cerevisiae* Rts1 residues 1-150 (Uniprot P38903) | Downey/Rudner labs |
| NN284 | Rts1(1-65) | N-MBP-TEV-insert-C | pET16b | *S. cerevisiae* Rts1 residues 1-65 (Uniprot P38903) | This study |
| NN140 | MafB(80-232) | N-MBP-TEV-insert-C | pET16b | Residues 80-232 of codon optimized MafB from NN145 template | This study |
| NN281 | MafB(80-232) ΔHis | N-MBP-TEV-insert-C | pET16b | As above, but with residues 131-167 deleted | This study |
| NN151 | Snf1 | N-MBP-TEV-insert-C | pET16b | *S. cerevisiae* WT Snf1 (Uniprot P06782) | Codon optimized for *E. coli* by GenScript |
| NN202 | Snf1 H>R | N-MBP-TEV-insert-C | pET16b | *S. cerevisiae* Snf1 with H18-H30 substituted to arginine residues | This study |
| NN167 | Snf1 ΔHis | N-MBP-TEV-insert-C | pET16b | *S. cerevisiae* Snf1 with residues 1-32 deleted | This study |
| KL4 | Snf1(1-65) WT | N-MBP-TEV-insert-C | pET28a | *S. cerevisiae* Snf1 WT residues 1-65 | Codon optimized for *E. coli* by Sangon Biotech |
| KL8 | Snf1(1-65) A1 | N-MBP-TEV-insert-C | pET28a | *S. cerevisiae* Snf1 residues 1-65 with H23A substitution | |
| KL9 | Snf1(1-65) A2 | N-MBP-TEV-insert-C | pET28a | *S. cerevisiae* Snf1 residues 1-65 with H22A and H27A substitutions | |
| KL23 | Snf1(1-65) A3 | N-MBP-TEV-insert-C | pMAL-C4X | *S. cerevisiae* Snf1 residues 1-65 with H21A, H22A, H26A, and H27A substitutions | Codon optimized for *E. coli* by GenScript |

| | | | | | |
|---|---|---|---|---|---|
| KL24 | Snf1(1-65) A4 | N-MBP-TEV-insert-C | pMAL-C4X | *S. cerevisiae* Snf1 residues 1-65 with H19A, H22A, H25A, H27A, and H30A substitutions | |
| KL25 | Snf1(1-65) A5 | N-MBP-TEV-insert-C | pMAL-C4X | *S. cerevisiae* Snf1 residues 1-65 with H20A, H22A, H25A, H27A, and H29A substitutions | |
| NN220 | DYRK1A IDR | N-MBP-TEV-insert-EGFP-C | pET16b | DYRK1A residues 491-686 from codon optimized template NN239 | This study |
| NN221 | DYRK1A IDR ΔHis | N-MBP-TEV-insert-EGFP-C | pET16b | As above, but with residues 599-619 deleted | This study |
| NN267 | Cyclin T1 IDR | N-MBP-TEV-insert-EGFP-C | pET16b | Cyclin T1 residues 462-654 from codon optimized template NN227 | This study |
| NN268 | Cyclin T1 IDR ΔHis | N-MBP-TEV-insert-EGFP-C | pET16b | As above, but with residues 517-528 deleted | This study |
| NN204 | N/A | N-insert-mCherry-C | pmCherry-N1 | Empty pmCherry-N1 | Jia lab |
| NN209 | PA PPK1 | N-insert-mCherry-C | pmCherry-N1 | *P. aeruginosa* PA14 *ppk1* gene fusion to mCherry | This study |
| NN189 | DYRK1A | N-EGFP-insert-C | pEGFP-C1 | Human cDNA-derived *DYRK1A* gene fusion to EGFP | Dr. Susana de la Luna |
| NN210 | DYRK1A ΔHis | N-EGFP-insert-C | pEGFP-C1 | As above but with residues 599-619 deleted | This study |
| NN1 | PA PPK1 | N-His-TEV-insert-C | pET28a | Codon optimized *P. aeruginosa ppk1* (Uniprot Q02EC2) | Dr. Francisco Chavez |
| NN29 | EC PPK1 | N-insert-His-C | pET28a | *E. coli ppk* (Uniprot P0A7B1) | Jia lab |
| N/A | SM PPK1 | N-insert-His-C | pET28a | *S. marcescens ppk1* (Uniprot Q93Q74) | Jia lab |
